# Explicit Knowledge Gates Expectation Suppression in the Motor System: Evidence from a TMS Oddball Paradigm

**DOI:** 10.64898/2026.04.27.721216

**Authors:** Monique E. Cost-Chretien, Reuben Rideaux, Dominic M. D. Tran

## Abstract

Models of predictive processing propose that the brain continuously generates predictions about incoming sensory input, updating an internal model of the environment through prediction errors when those predictions are violated. A foundational assumption of these models is that prediction errors are generated automatically, even in the absence of explicit knowledge of the predictions. Evidence from auditory oddball studies in unconscious patients appears to support this view, though findings are complicated by stimulus-specific adaptation confounds that make it difficult to isolate genuine predictive effects. To investigate whether expectation suppression or prediction-based attenuation extends to the motor system and whether it operates automatically, we developed a novel motor oddball paradigm using brain stimulation. Transcranial magnetic stimulation (TMS) delivered over the primary motor cortex elicits motor-evoked potentials (MEPs) in peripheral muscles, providing an index of corticospinal excitability. By varying stimulation intensity in an oddball-like manner using repeating and deviating sequences, we manipulated the predictability of TMS pulses and compared MEP amplitudes for expected versus unexpected intensity-matched stimulation. Incorporating experimental designs to control for adaptation and an instruction manipulation to test the role of explicit knowledge, expected TMS produced smaller MEPs than unexpected TMS. Critically, this attenuation was observed only in participants explicitly informed about the sequence structure. These findings extend expectation suppression effects to the motor system and support the domain-generality of prediction-based neural attenuation while challenging the assumption that predictive processing operates entirely automatically.

## Introduction

The brain does not passively record the world — it actively predicts it. A central principle of modern neuroscience posits that the brain operates as a predictive machine, continuously constructing internal models of the environment based on prior experience and using these models to anticipate incoming sensory information (see Clark, 2013; Knill & Pouget, 2004, for reviews). When predictions are accurate, neural responses to expected inputs are suppressed; when predictions are violated, prediction errors signal the need to update the model (e.g., Friston, 2005; Lee & Mumford, 2003; Rao & Ballard, 1999). Theories of predictive processing have reshaped our understanding of perception, learning, and action.

Repetition-based paradigms such as the auditory oddball task have been central to studying predictive processes and neural attenuation in the laboratory. In these designs, a stream of repeated standard tones is occasionally interrupted by a deviant. The deviant reliably elicits a mismatch negativity (MMN) signal, a well-characterised EEG component reflecting a prediction error signal (den Ouden et al., 2012; Garrido et al., 2009). Crucially, Lieder et al. (2013) showed that trial-by-trial changes in MMN amplitude track updates to an internal probabilistic model rather than simple neural adaptation, establishing the MMN as an index of statistical learning. Analogous prediction error signalling and prediction-based attenuation (also commonly referred to as expectation suppression) effects have been documented in dopaminergic (Schultz et al., 1997), limbic (Tzovara et al., 2024), and somatosensory (Akatsuka et al., 2007) systems, pointing to the possibility that predictive processes are a domain-general computational principle (Tran & Livesey, 2021).

More recently, expectation suppression has also been demonstrated in the motor system. When transcranial magnetic stimulation (TMS) is delivered over primary motor cortex and can be anticipated, either through self-generation, visual cueing, or temporal regularity, the resulting motor-evoked potentials (MEPs) are suppressed relative to MEPs elicited by unexpected pulses of identical intensity (Sriutaisuk & Franz, 2025; Tran et al., 2021). Using combined TMS-EEG, Tran et al. (2025) further showed that the magnitude of motor attenuation correlated with the magnitude of sensory attenuation measured concurrently to the TMS coil click, suggesting a shared predictive mechanism across motor and sensory circuits.

A foundational assumption of predictive processing accounts is that the brain continuously operates, automatically computing prediction errors at every level of the cortical hierarchy, and largely independently of conscious awareness or voluntary control (Clark, 2013; Friston, 2005; Hutchinson & Barrett, 2019). Evidence for this hypothesis comes from auditory oddball studies conducted in unconscious participants (sleeping, anaesthetised, or in a disorder of consciousness), which demonstrate that MMN-like prediction error responses can survive the loss of consciousness (Fischer et al., 1999; Koelsch et al., 2006; Simpson et al., 2002), and that their presence can even predict recovery from coma (Tivadar et al., 2021). These findings have been widely interpreted as evidence that predictive processes operate independently of awareness or explicit knowledge of the deviance (unexpected) structure (Tivadar et al., 2021). However, this interpretation has been strongly contested: when stimulus-specific adaptation is controlled using expectation designs that vary the frequency of local and global deviants, evidence for automatic global expectation effects in unconscious patients is substantially reduced (Bekinschtein et al., 2009; Nourski et al., 2018; Strauss et al., 2015), challenging the automaticity assumption.

Converging evidence that challenges the automatic hypothesis comes from designs where predictions are established through learned associations rather than repetition structure. In the visual domain, Meijs et al. (2018) and Alilović et al. (2021) used an attentional blink paradigm and found that a valid prediction facilitated conscious access to a target only when the predictive cue was itself consciously perceived. Additionally, the benefit scaled with explicit knowledge of the contingency between the predictive cue and the target (Meijs et al., 2018). In the motor domain, the role of contingency awareness and explicit knowledge was tested by Liang et al. (2025) using a masking paradigm in which a visual stimulus covertly predicted an upcoming TMS pulse across two interleaved tasks. Motor attenuation was observed only in participants who were explicitly informed of the predictive relationship between the two tasks or who spontaneously became aware of it; participants who remained unaware showed no attenuation. Together, these findings challenge the assumption that prediction error signals are generated automatically.

A limitation of the Liang et al. (2025) design, however, is that predictability was cued cross-modally, with visual stimuli signalling motor stimulation. Cross-modal integration imposes additional cognitive demands (Sepulcre et al., 2012) and may itself require awareness, potentially masking automatic processes that might otherwise emerge. Sensory paradigms avoid this confound by cueing predictability within a single modality (Rideaux et al., 2025). In auditory oddballs, both the standard sequence that establishes the expectation and the deviant target that violates the expectation are tones. The same is true in visual cueing paradigms such as the attentional blink, where both the predictive cue and the target are visual stimuli (e.g., letters). This unimodal architecture arguably provides a stronger test of whether automatic predictive processes exist.

Here, we addressed this gap within the motor prediction literature by developing a novel TMS-based motor oddball paradigm that mirrors the unimodal structure of sensory paradigms. The ingenuity here is that expectation and violations of expectation are produced and measured entirely by stimulating the motor cortex. Across three experiments, we tested the effectiveness of TMS motor oddball paradigm and asked whether expectation suppression could be elicited automatically or whether, even under optimal conditions, explicit knowledge of the predictive structure remains necessary. Experiment 1 established that TMS can produce repetition-based motor attenuation under sufficiently high stimulation intensities. Experiment 2 implemented a local-global expectation design to dissociate genuine predictive effects from stimulus-specific adaptation. Experiment 3 tested whether explicit knowledge of oddball sequence structure is required for motor attenuation. Converging across experiments, our results demonstrate that a TMS oddball design can elicit expectation suppression. Furthermore, this form of prediction-based attenuation depended on explicit knowledge of oddball sequence, adding to a growing body of evidence challenging accounts that assume predictive processing operates automatically.

## Results

### Experiment 1

Predictability was manipulated by varying the stimulation intensity of TMS (see Figure 1A). Repeat sequences of low intensity and high intensity pulses (i.e., low, low, low, low; and high, high, high, high) comprised the majority of trials and established an expectation of repeating intensity stimulation. Deviant sequences (i.e., low, low, low, high; and high, high, high, low) comprised the minority of trials and violated the expectation of repeats. Given well-documented variability in raw MEP amplitudes between participants, primary analyses used log-normalised MEPs (see Tran et al., 2021, for detailed discussion on best practice analysis) of expected repeat and unexpected deviant sequences to enable rescaled comparisons between participants; negative values indicate smaller MEP for the expected relative to unexpected conditions (i.e., expectation suppression). The factorial analyses of the raw data (Figure 1B) are presented in the supplementary materials. Overall, collapsing across intensity, there was no significant expectation suppression effect, *t*(28) = –1.46, *p* = .156, *d* = –.271 (Figure 1C).

**Figure 1.**
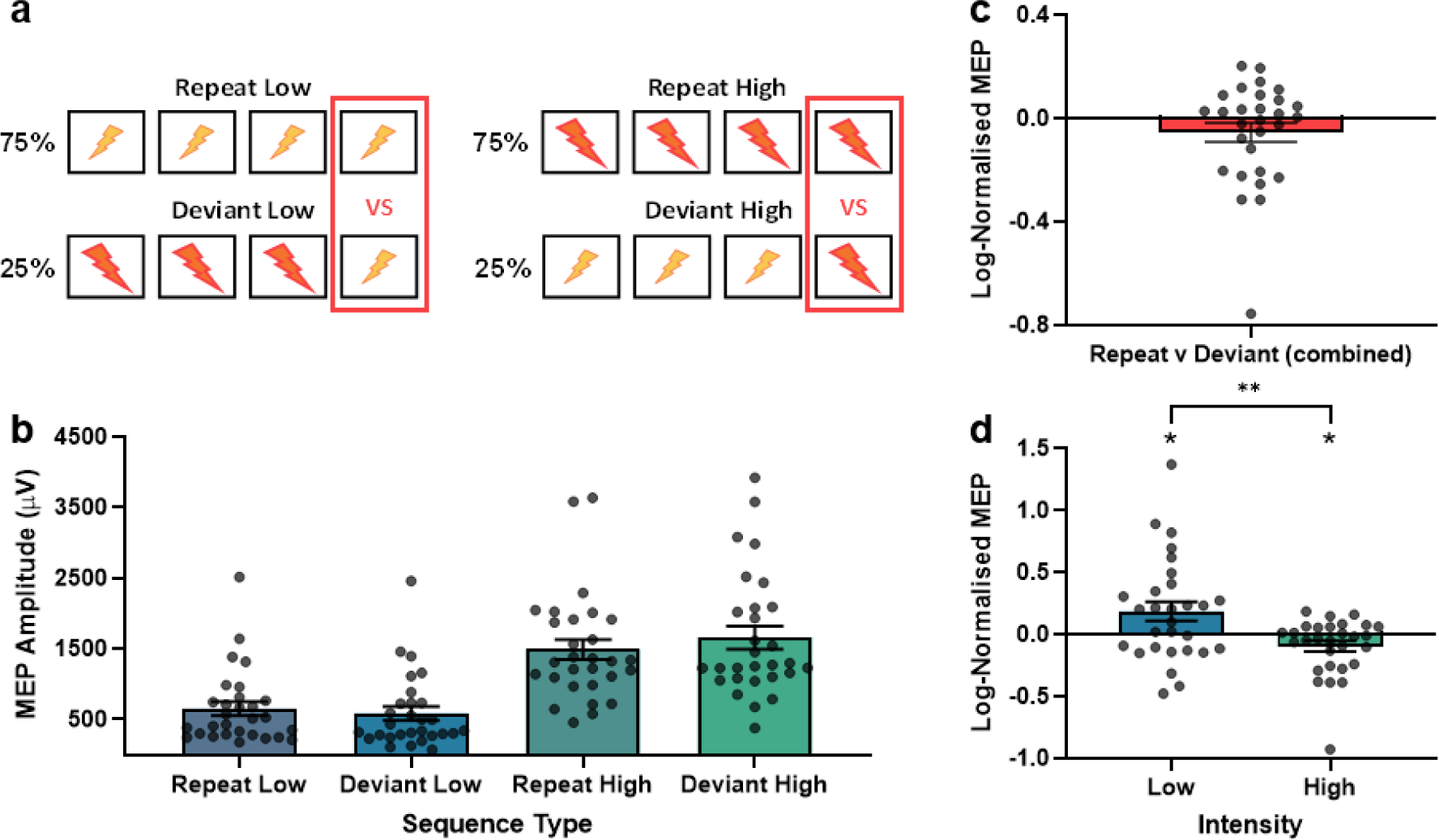
Schematic design and results for Experiment 1. A) Schematic representation of oddball sequence types. B) Mean raw MEP amplitudes by sequence type and stimulation intensity. C) Mean log-normalised MEPs for Repeat (expected) sequences relative to Deviant (unexpected) sequences collapsed across stimulation intensity. D) Mean log-normalised MEPs of Repeat (expected) sequences relative to Deviant (unexpected) sequences separated by stimulation intensity. *Note*. For log-normalised MEPs, negative values indicate attenuation of expected pulses relative to unexpected pulses, and positive values indicate elevation of expected pulses relative to unexpected pulses. Error bars represent standard error of the mean. * p < .05 ** p < .01

However, splitting by intensity; there was significant attenuation for high intensity stimulation (deviant high versus repeat high), *t*(28) = –2.21, *p* = .036, *d* = –.409, and significant elevation for low intensity stimulation (deviant low versus repeat low), *t*(28) = 2.42, *p* = .022, *d* = .450 (see Figure 1D). The difference in attenuation between intensities was also significant, *t*(28) = 2.89, *p* = .007, *d* = .537. The absence of motor attenuation for low intensity stimulation washed out the overall expectation suppression effect when collapsing across intensity, and likely reflects stimulus-specific adaptation: repeated exposure to high intensity pulses in deviant low sequences (high, high, high, low) reduces sensitivity to the final low intensity pulse, obscuring any excitability change relative to repeat low sequences (low, low, low, low). This pattern mirrors intensity-dependent adaptation effects in visual perception, where prior exposure to a bright stimulus impairs detection of a subsequent dim one (Normann & Perlman, 1979; Stevens & Stevens, 1963). These results demonstrate that expected brain stimulation produces attenuated motor responses relative to unexpected stimulation, provided that the stimulation intensity is sufficient to detect such effects (deviant high versus repeat high).

### Experiment 2

The motor attenuation to repeat high sequences (high, high, high, high) from Experiment 1 may be driven by expectation of high repeating pulses; within sequence (local) adaptation following repeated high intensity stimulation; or a combination of both. To control for possible adaptation, Experiment 2 manipulated predictability at both local and global levels by varying the frequency of repeat and deviant sequences across blocks (Figure 2). In global repeat blocks, repeat sequences comprised the majority of trials and deviant sequences the minority, replicating Experiment 1. In global deviant blocks, these frequencies were reversed: deviant sequences were predominant, establishing an expectation of deviance, while repeat sequences constituted the minority and therefore violated that expectation.

**Figure 2.**
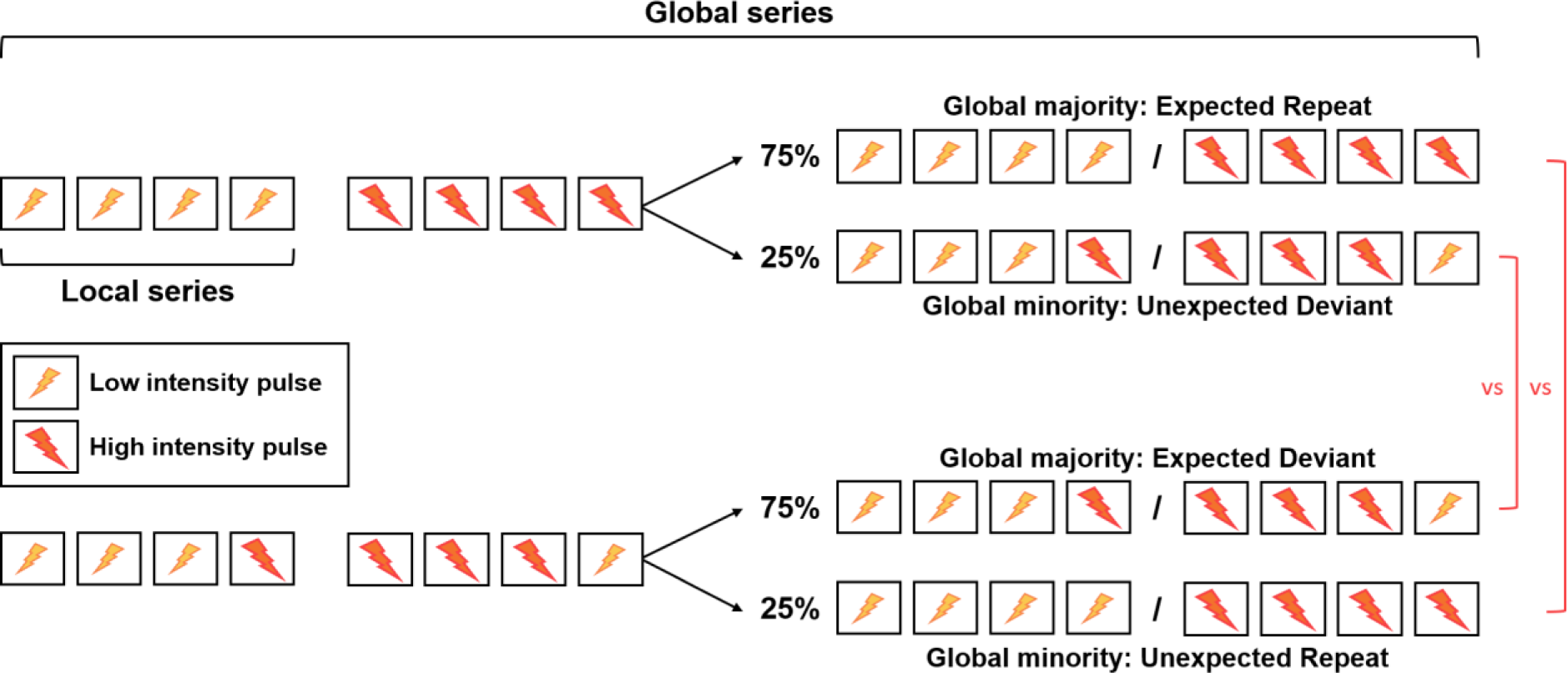
Schematic representation of the local-global design used in Experiment 2. *Note*. Global series refers to the block level, and local series refers to the trial level. Two types of global series were used: one in which Repeat trials were the global majority, and one in which Deviant trials were the global majority. Majority trials occurred 75% of the time and minority trials occurred 25% of the time.

The factorial analyses of the raw data (Figure 3A) are presented in the supplementary materials. Log-normalised MEPs of global majority (expected) to global minority (unexpected) sequences controlling for between-participant variability revealed a significant overall prediction effect, with expected TMS producing smaller MEPs than unexpected TMS, *t*(23) = –2.33, *p* = .029, *d* = – 0.475 (Figure 3B). These findings confirm that a TMS-based oddball paradigm is sufficiently sensitive to detect expectation suppression in the motor system when adaptation and between-participant variability are adequately controlled. Split by sequence type, there was significant attenuation for deviant sequences, *t*(23) = –4.78, *p* < .001, *d* = –0.975, but not for repeat sequences, *t*(23) = 1.89, *p* = .071, *d* = 0.386. The difference in attenuation between sequence type was also significant, *t*(23) = 4.17, *p* < .001, *d* = 0.850 (see Figure 3C). This asymmetry in attenuation for deviant versus repeat sequences may reflect local adaptation exerting a stronger influence on motor excitability than global frequency for repeat sequences. One possible account is that the expectation of a fourth high-intensity pulse in deviant majority blocks (low, low, low, high) shifts the stimulus-response function, reducing sensitivity to the final low-intensity pulse in unexpected repeat sequences (low, low, low, low). That is, expecting a high pulse makes a low pulse hard to detect, obscuring any relative increase in excitability due to being unexpected compared to the same sequence type when it is expected in repeat majority blocks. Prior studies using local-global designs have typically reported overall prediction effects collapsed across sequence types, a result replicated here. However, sequence-level breakdowns are seldom reported (e.g., Summerfield et al., 2008; Tang et al., 2018), making it difficult to determine whether the differential sensitivity between deviant and repeat sequences observed here is characteristic of this paradigm more broadly.

**Figure 3.**
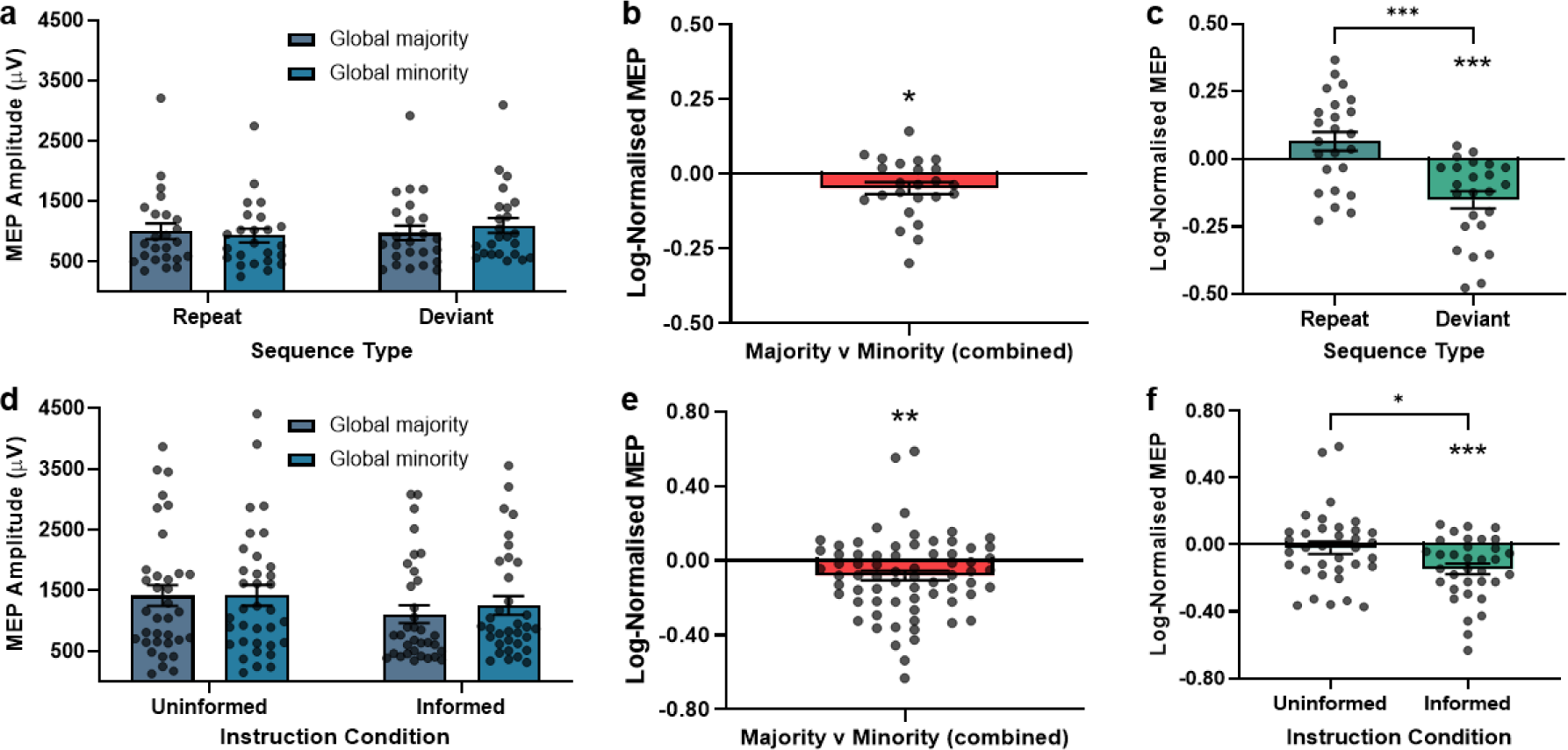
Results for Experiments 2 (top row: A-C) and 3 (bottom row: D-F). A) Mean raw MEP amplitudes by sequence type and predictability. B) Mean log-normalised MEPs for global majority (expected) sequences relative to global minority (unexpected) sequences, collapsed across sequence type (repeat and deviant sequences). C) Mean log-normalised MEPs for global majority (expected) sequences relative to global minority (unexpected) sequences separated by sequence type. D) Mean raw MEP amplitudes by instruction condition and predictability. E) Mean log-normalised MEPs for global majority (expected) sequences relative to global minority (unexpected) sequences, collapsed across instruction condition (uninformed and informed) F) Mean log-normalised MEPs for global majority (expected) sequences relative to global minority (unexpected) sequences by instruction condition. *Note.* For log-normalised MEPs, negative values indicate attenuation of expected pulses relative to unexpected pulses. Error bars represent standard error of the mean. * p < .05 ** p < .01 *** p < .001

### Experiment 3

Experiment 3 examined whether knowledge of the trial and block structures is necessary for expectation suppression. Global frequency was manipulated within participants as in Experiment 2, and explicit knowledge was manipulated between participants: the Informed group received instructions about both the local sequence structures and the global block frequencies (same as Experiment 2), whereas the Uninformed group received minimal instructions that TMS pulses would occur in succession. If prediction errors are generated automatically, prediction-based attenuation should be observed regardless of instruction condition (Tivadar et al., 2021); if explicit knowledge of the task structure is required, motor attenuation should be restricted to the Informed group (Bekinschtein et al., 2009; Liang et al., 2025; Meijs et al., 2018). Given the sensitivity limitations identified for low intensity and repeat sequences in Experiments 1 and 2, primary analyses focused on deviant sequences, which produced the most reliable expectation suppression effects (see Methods for full details).

The factorial analysis of the raw data (Figure 3D) are presented in the supplementary materials. Log-normalised analyses found a significant overall prediction effect, with expected TMS producing smaller MEPs than unexpected TMS, *t(68) =* –3.16*, p =* .002*, d = –*.381 (Figure 3E). Follow-up analyses revealed there was significant attenuation in the Informed group, *t*(33) = – 4.51, *p* < .001, *d* = –.774, but not in the Uninformed group, *t*(34) = –0.51, *p* = .614, *d* = –.086. The difference in attenuation between groups was also significant, *t*(67) = 2.56, *p* = .013, *d* = .615 (Figure 3F). These results indicate that explicit knowledge of both local and global sequence structures is necessary for the emergence of expectation suppression in the motor system.

## Discussion

The present study investigated expectation suppression in the motor system using a novel TMS oddball paradigm. Across three experiments, expected TMS pulses consistently elicited smaller MEPs than unexpected pulses that were matched for intensity. The results replicate previous demonstrations of expectation suppression in the motor domain using self-generated and visually cued prediction (Liang et al., 2025; Sriutaisuk & Franz, 2025; Tran et al., 2021; Tran et al., 2025) and extend these findings, for the first time, to repetition-based prediction. Critically, expectation suppression remained significant when stimulus-specific adaptation was controlled using a local-global design, confirming that predictive processes independently contribute to motor attenuation beyond adaptation alone.

### Sensitivity of the TMS oddball paradigm

Although the overall pattern of results supports the validity of the TMS oddball approach, the sensitivity of the paradigm varied across sequence types. In Experiment 1, expectation suppression was detected for high intensity but not low intensity stimulation. This asymmetry likely reflects intensity-dependent adaptation. In Experiment 2, a similar pattern emerged, with reliable expectation suppression observed for deviant but not repeat sequences. The absence of motor attenuation for repeat sequences may reflect a shift in the stimulus-response function. Across all experiments, deviant high intensity sequences proved to be the most reliable index of expectation suppression. Prior studies using local-global designs have typically reported overall prediction effects collapsed across sequence types without separating deviant and repeat sequences (e.g., Summerfield et al., 2008; Tang et al., 2018), making it difficult to determine whether differential sensitivity between sequence types is a general feature of such designs. Future work should examine whether analogous sensitivity differences emerge in auditory oddball paradigms and investigate the mechanisms underlying this asymmetry.

### The role of explicit knowledge in expectation suppression

Experiment 3 tested whether explicit knowledge of the trial and block structure is necessary for expectation suppression using a between-participant manipulation. Participants who were informed about both local trial sequences and global block frequencies showed robust motor attenuation, whereas uninformed participants did not. The results from the informed group of Experiment 3 also replicate the findings of Experiment 2, which essentially mimicked a single informed group design using similar instructions. This pattern is consistent with Liang et al. (2025), who found that motor attenuation was restricted to participants who were explicitly informed of, or who spontaneously became aware of, a predictive relationship between visual cues and TMS pulses (see also Meijs et al., 2018). Together, these findings suggest that expectation suppression in the motor system is not generated entirely automatically but depends on explicit knowledge of the task structure and conscious awareness of the predictive contingency.

This interpretation also aligns with recent evidence showing that predictive processing tracks the represented (subjective) rather than the veridical (objective) statistical structure of a task. Clarke et al. (2026) had participants discriminate probabilistically cued visual outcomes in addition to reporting post-experiment probability, trial-by-trial expectancy, or trial-by-trial surprise associated with each outcome. They found that the subjective reports explained variance in reaction time and accuracy over and above the objective contingencies, with perceptual accuracy in some cases best explained by the subjective ratings alone. The findings by Clarke et al. (2026) characterise the contribution of subjective predictions as graded and additive, among participants who were all aware of the task structure. The present results, however, speak to a more fundamental question: whether predictive processes arise at all in the absence of explicit knowledge of the task structure. Based on the available evidence, explicit knowledge of the predictions is required for expectation suppression (Meijs et al., 2018), and the current results extend this to the motor system.

An important consideration for Experiment 3 is that while a local-global design controlled for local adaptation effects, using only half of the sequence types from Experiment 2 may have introduced differences in global adaptation (see Methods). Specifically, Deviant common blocks contained a greater number of high intensity pulses compared to Repeat common blocks. However, the results from Experiment 2, which employed a fully balanced design, demonstrate that prediction and expectation produce an observable and equivalent-sized effect over and above adaptation in this TMS oddball paradigm. Therefore, although some portion of the attenuation observed in Experiment 3 may reflect adaptation, it is unlikely that the attenuation is driven entirely by adaptation and likely that predictive mechanisms contribute to the effect.

We had intended to complement the instruction manipulation with an assessment of awareness derived from a post-experimental questionnaire. However, the unique phenomenological experience of TMS made it difficult for participants to reliably report variations in stimulation intensity. For example, some participants reported experiencing more than the two (low and high) intensities delivered, even in participants who were explicitly informed about the trial structures. This result precluded a valid assessment of their knowledge of the sequence structure and therefore an aware versus unaware classification based on the questionnaire was deemed to be invalid (see Methods and Supplemental Materials for full discussion).. Future research could circumvent this issue by studying participants in various awareness states, or by developing more sensitive awareness measures tailored to intensity-based paradigms The current findings stand in apparent contrast to studies reporting that sensory prediction error signals, such as the MMN, can be elicited in unconscious patients who are presumably unaware of the predictive relationships (Fischer et al., 1999; Koelsch et al., 2006; Simpson et al., 2002). However, the traditional oddball paradigms used in those studies are inherently confounded by stimulus-specific adaptation, raising the possibility that the observed effects reflect adaptation rather than genuine predictive processing (Feuerriegel et al., 2021).

Consistent with this concern, studies that employ local-global designs to control for adaptation have generally failed to find global prediction effects in unconscious or mind-wandering participants (Bekinschtein et al., 2009; Nourski et al., 2018; Strauss et al., 2015). When considered alongside the present findings, this body of evidence challenges the assumption that predictive model updating in response to unexpected information occurs entirely automatically (Tivadar et al., 2021).

### Reconsidering the MMN as a marker of automatic prediction

A widely cited line of evidence for automatic predictive processing is the finding that MMN amplitude predicts recovery from coma, which has been interpreted as evidence that prediction error signals can emerge in the absence of explicit knowledge of the predictions (Naccache et al., 2005; Pfeiffer et al., 2018; Tivadar et al., 2021). However, this interpretation warrants scrutiny. Patients in a shallower coma state are generally more physiologically responsive and more likely to recover consciousness (Schnakers & Laureys, 2012); they may also, for the same reasons, produce stronger MMN responses. On this account, the MMN-recovery correlation need not imply that prediction errors are generated without explicit knowledge — it may simply reflect that both outcomes are jointly predicted by coma depth. Stronger evidence would require demonstrating that MMN amplitude predicts recovery after controlling for coma depth. Neither Naccache et al. (2005) nor Pfeiffer et al. (2018) performed this control: the former restricted their sample to patients with a Glasgow Coma Scale score below 8, and the latter focused on post-anoxic comatose patients, but neither study statistically accounted for coma depth when relating MMN to outcome. These findings therefore do not constitute strong evidence that prediction errors manifest automatically in the absence of explicit knowledge.

### A domain-general predictive mechanism and clinical implications

The present results also speak to broader questions about the architecture of predictive processing in the brain. Previous work has demonstrated expectation suppression when TMS predictability is established via visual or temporal cues (Tran et al., 2021); the current study extends this to repetition-based prediction, bringing the motor paradigm into closer alignment with the sensory literature (Elijah et al., 2018; Tivadar et al., 2021). Moreover, Tran et al. (2025) showed that individual differences in sensory attenuation predicted individual differences in motor attenuation, suggesting that the degree to which predictive processes modulate neural responses may reflect a domain-general trait that varies across individuals. This finding points toward a unified predictive mechanism operating across sensory and motor systems rather than being independent modality-specific processes.

Such individual variability in predictive processing may have important clinical implications. A growing body of work implicates atypical prediction error signalling in conditions including schizophrenia, autism spectrum disorder, and chronic pain (see Tschentscher et al., 2023; van Schalkwyk et al., 2017, for reviews). The TMS oddball paradigm introduced here offers a novel tool for quantifying predictive processing in the motor system that could complement existing sensory measures in clinical populations, potentially providing a more comprehensive characterisation of individual differences in predictive processing across domains.

### Conclusion

The present study introduces a TMS oddball paradigm as a new tool for studying predictive processes in the motor system and provides the first demonstration of repetition-based motor attenuation. Across three experiments, expected brain stimulation produced attenuated motor responses relative to unexpected stimulation, an effect that persisted after controlling for stimulus-specific adaptation. Critically, attenuation was restricted to participants with explicit knowledge of the predictive structure, supporting the view that expectation suppression in the motor system is not entirely automatic. These findings contribute to a growing body of evidence challenging the automaticity hypothesis within predictive processing accounts and highlight the utility of unimodal, repetition-based paradigms for probing the boundaries of automatic and explicit knowledge of predictions across neural systems.

## Methods

### Design and Procedure

All experiments employed a passive oddball task that did not require participants to engage in a secondary task (e.g., counting or detection). The absence of a secondary task allowed participants to become aware of the sequence regularities, since distraction can interfere with global expectations (Bekinschtein et al., 2009). Moreover, it was important that participants keep their hand at rest for the MEP recordings. However, given the passive nature of the task, to ensure participants did not mind wander, they were instructed to pay close attention during the task: “You do not need to do anything during the experiment but please try and stay attentive because we will ask you some simple questions at the end of the experiment.” The instruction was emphasised verbally by the experimenter, and breaks were included after each block to help maintain focus.

#### Experiment 1

The design mirrored a classic auditory oddball task, adapted for TMS intensity. In standard auditory oddball tasks, stimuli are delivered in a rapid continuous stream with approximately 500 ms between stimulus onsets (Koshiyama et al., 2020). Because TMS equipment requires approximately four seconds to recharge between pulses, two stimulators (Unit A and Unit B) were programmed to deliver pulses in alternation (A, B, A, B), producing a 2-second interstimulus interval. Each trial therefore comprised a sequence of four TMS pulses.

Trials were arranged in a 2 (Intensity: low, high) × 2 (Predictability: expected, unexpected) design. Repeat sequences (low, low, low, low; high, high, high, high) comprised the majority of trials and established an expectation of repeating intensity. Deviant sequences (low, low, low, high; high, high, high, low) comprised the minority of trials and violated that expectation on the final pulse. This produced four sequence types: Repeat Low, Repeat High, Deviant Low, and Deviant High (see Figure 1).

Across five blocks, sequence predictability was established by presenting Repeat sequences on 75% of trials and Deviant sequences on the remaining 25%. Within each block, trials were pseudorandomised into two sets of 16 (12 Repeat and 4 Deviant). At the beginning of the first block, one additional Repeat High and one additional Repeat Low trial were presented in random order to establish an expectation of repeating sequences prior to data collection; these trials were excluded from analysis. This yielded 160 trials (5 blocks × 2 sets × 16 trials) for analysis.

One potential confound is that the TMS device produces a faint auditory artefact when switching from high to low stimulation intensity, due to the discharge of stored electrical charge. This artefact occurs during Deviant Low sequences (high, high, high, low) and after trials ending with a high pulse, as the device was programmed to reset to low intensity at the end of each sequence. If detected, this sound could serve as a cue signalling an upcoming Deviant Low pulse. To minimise this confound, participants wore earplugs throughout the experiment. A post-experimental debrief confirmed that no participant reported detecting this auditory artefact.

#### Experiment 2

The design followed a local-global paradigm in which the same sequence types varied in their global predictability across blocks according to their relative frequency (see Figure 2). Repeat sequences (Repeat Low, Repeat High) and Deviant sequences (Deviant Low, Deviant High) each alternated between serving as the global majority (expected; 75%) and the global minority (unexpected; 25%) across 10 blocks. The sequence type assigned to the majority on the first block was counterbalanced across participants.

Each block began with two trials of the globally expected sequence type (one of each stimulation intensity) to establish the global expectation; these trials were excluded from analysis. The remaining trials were pseudorandomised into four sets of eight (6 expected and 2 unexpected), yielding 320 trials (10 blocks × 4 sets × 8 trials) for analysis.

Prior to commencing the experiment, participants were informed about the local-global structure of the task, including the different sequence types and how their frequency varied across blocks. At the start of each block, participants were informed which sequence type would occur most frequently.

#### Experiment 3

The design used the same local-global paradigm as Experiment 2 but included only Repeat Low and Deviant High sequences. This decision was made because the Deviant Low condition had a potential auditory confound (described in Design and Procedure – Experiment 1), limited sensitivity following high intensity pulse (described in Results – Experiment 1), and for practical considerations due to the longer time required to run the full design compounded with the additional between-participant manipulation. Deviant High sequences were selected because they produced the most reliable and largest expectation suppression effects across Experiments 1 and 2; Repeat Low sequences were retained as their counterpart. Given the sensitivity limitations identified for Repeat Low sequences in Experiment 2, however, primary analyses focused on comparing expected Deviant High sequences with unexpected Deviant High sequences.

Across six blocks, Deviant High and Repeat Low sequences alternated between serving as the global majority (75%) and global minority (25%). Each block began with two trials of the globally expected sequence type to establish the expectation; these trials were excluded from analysis. The remaining trials were pseudorandomised into four sets of eight (6 expected and 2 unexpected), yielding 192 trials (6 blocks × 4 sets × 8 trials) for analysis.

Participants were randomly assigned to one of two instruction groups. The Informed group received the same briefing as participants in Experiment 2, including information about the different sequence types and which sequence would occur most frequently at the start of each block. The Uninformed group were simply instructed that TMS pulses would be delivered in succession.

To verify engagement with the task, participants in the Informed group completed a manipulation check after each block, selecting which sequence had occurred most frequently: “Sequence 1 (low, low, low, low)” or “Sequence 2 (low, low, low, high)”. On average, participants correctly identified the majority sequence on 93.8% of blocks, and no participant answered more than half of the checks incorrectly; all participants were therefore retained for analysis. To match the timing and cognitive demands between groups, participants in the Uninformed group completed an analogous filler question after each block, reporting how much their head had moved during stimulation.

Participants in both groups also completed a post-experimental questionnaire designed to assess awareness of the sequence structure. The questionnaire was adapted from previous work assessing awareness in visual cueing tasks (Liang et al., 2025; Weidemann et al., 2016). However, accurately probing awareness in the context of a TMS intensity paradigm proved difficult for several reasons. The experience of TMS stimulation is entirely novel and unlike other sensory or motor events participants have previously encountered, making it challenging to accurately probe phenomenological experience through self-report. Additionally, expected and unexpected TMS pulses of identical physical intensity may nonetheless be experienced differently, for instance, unexpected pulses may feel stronger. This effect would be consistent with reports of expectation-dependent perceptual differences in other modalities (Dewey & Carr, 2013; Eagleman & Pariyadath, 2009). As a result, the questionnaire lacked the sensitivity and validity required to classify participants as aware or unaware of the sequence structure, and the primary awareness analysis therefore relied on the instruction group manipulation. The full questionnaire including a detailed discussion of its limitations and the problems encountered are provided in the Supplemental Materials.

### Participants

Participants were first- and second-year psychology students who received course credit for participation. All participants completed a TMS safety screening questionnaire (Rossi et al., 2021) and provided written informed consent prior to commencing the experiment. All procedures were approved by the Human Research Ethics Committee of The University of Sydney.

As this was the first series of studies to investigate expectation suppression using a TMS oddball paradigm, no comparable effect size was available to inform an a priori power analysis. Target sample sizes were therefore guided by previous TMS studies of prediction-based motor attenuation, which have typically employed samples of 24–30 participants (Tran et al., 2021).

#### Experiment 1

We aimed to recruit 30 participants. Thirty-three participants were initially enrolled; three were excluded due to a resting motor threshold (rMT) exceeding 70% of maximum stimulator output (MSO), and one was excluded because the thickness of their head wrap resulted in insufficient proximity between the TMS coil and the scalp. To allow differentiation between low and high stimulation intensities, pulse intensities were set at 110% of rMT rather than the 120% typically used in the laboratory (see Table 1). Because lower stimulation intensities can produce smaller MEPs, participants were additionally excluded if their mean MEP amplitude fell below 250 μV; no participants met this criterion. The final sample comprised 29 participants (mean rMT = 49.41% MSO, SD = 7.54).

**Table 1.** Low and High Intensity Configurations.

| rMT | Low | High |
| --- | --- | --- |
| $\leq 45$ | 50% MSO | 62% MSO |
| $45 < \text{rMT} < 55$ | 110%*rMT | 135%*rMT |
| $55 < \text{rMT} < 65$ | 110%*rMT | 130%*rMT |
| $\geq 65$ | 110%*rMT | 125%*rMT |

*Note.* MSO = Maximum Stimulator Output

#### Experiment 2

We again aimed to recruit 30 participants. Thirty-three participants were initially enrolled; four were excluded due to an rMT exceeding 70% MSO. A further five were excluded prior to analysis: one required a translator to understand the task instructions, one fell asleep during the experiment, and three exhibited excessive head movement that resulted in a mean MEP amplitude below 250 μV. The final sample comprised 24 participants (mean rMT = 52.25% MSO, SD = 7.21).

#### Experiment 3

Given the between-subjects comparison, we aimed to recruit 80 participants across two equal groups (40 Informed, 40 Uninformed). Eighty-one participants were initially enrolled; one was excluded due to an rMT exceeding 70% MSO. A further 11 were excluded prior to analysis: five due to difficulty identifying a stable TMS hotspot, two due to excessive head movement compromising coil placement and their MEP recording, two who fell asleep during the session, and two who exhibited large and highly variable hand twitches that compromised data quality. The final sample comprised 69 participants — 34 in the Informed condition and 35 in the Uninformed condition (mean rMT = 49.49% MSO, SD = 8.22).

### Stimuli and Apparatus

#### Electromyography (EMG)

Surface EMG activity was recorded from the first dorsal interosseous (FDI) muscle of the right hand using a pair of Ag/AgCl electrodes placed over the proximal and distal muscle belly, with a ground electrode positioned over the ulnar styloid process of the wrist. EMG was recorded from 200 ms prior to TMS delivery to 100 ms following stimulation using a PowerLab 26T data acquisition device (ADInstruments, Bella Vista, NSW, Australia). Signals were digitised at a sampling rate of 4 kHz, bandpass filtered between 0.5 Hz and 2 kHz, and notch filtered at 50 Hz (mains), then stored for offline analysis using LabChart software (Version 8, ADInstruments).

#### Transcranial Magnetic Stimulation (TMS)

TMS was delivered using two Magstim 2002 stimulators connected to a 70-mm figure-of-eight coil. A cotton cap was secured to the participant’s head to assist with coil localisation over the left primary motor cortex (M1). The motor hotspot, defined as the scalp location eliciting the largest and most consistent MEPs in the FDI muscle, was identified by positioning the coil approximately 5 cm lateral and 1 cm anterior to Cz and then systematically adjusting its position. Once identified, the coil was fixed tangentially over the hotspot with the handle oriented at approximately 45° from the midsagittal axis using a Manfrotto articulating arm. An adjustable forehead and chin rest was used to minimise head movement throughout the experiment.

Resting motor threshold (rMT) was determined for each participant prior to the oddball task, defined as the lowest stimulator output capable of inducing MEPs with a peak-to-peak amplitude of at least 50 μV in at least 50% of consecutive pulses (Rossini et al., 2015). Low and high pulse intensities for the oddball task were then set individually based on each participant’s rMT. Intensities were determined through pilot testing to ensure that the low intensity pulse was suprathreshold, that the two intensities were perceptually discriminable, and that the high intensity remained within a comfortable range for participants (see Table 1).

#### Task Stimuli

The experiment was run on a PC using Python to present stimuli and collect response data. This was connected to a 24-inch ASUS monitor (1920 x 1080 resolution, 60 Hz refresh rate) at a viewing distance of approximately 63 cm.

### Data Analysis

MEP data were pre-processed using custom software written in Python. Trials were excluded if background EMG activity exceeded 100 μV peak-to-peak amplitude during the pre-stimulation interval, or if EMG activity exceeded 100 μV root mean square over the entire 200 ms pre-stimulation period. For retained trials, peak-to-peak MEP amplitude was extracted as the difference between the maximum and minimum EMG values in the post-stimulus window, prior to signal return to baseline.

Additional trial-level exclusions were applied when the TMS device disarmed mid-trial or at the end of a block due to stimulator constraints (e.g., overheating). Trials and blocks were also excluded when excessive participant movement or coil displacement from the hotspot was observed and documented. Such instances were identified by cross-referencing block-level indicators, including mean MEP amplitude below 250 μV and a high number of zero-amplitude MEP trials (> 10 per block). In Experiment 3, however, the 250 μV block-level exclusion criterion was not applied, as the high frequency of low intensity stimulation at 110% rMT in Repeat Low majority blocks would have resulted in disproportionate exclusion of those blocks. This criterion was similarly not applied at the participant level in Experiment 3 for the same reason. Across all three experiments, fewer than 10% of trials were excluded in total.

Mean MEP amplitudes were log-normalised at the participant level to account for large individual differences in absolute MEP amplitude and to correct for positively skewed distributions (Tran et al., 2021). Log-normalised values significantly less than zero indicate attenuation of expected relative to unexpected sequences, as follows:

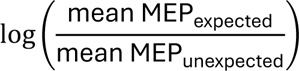

#### Experiment 1

A one-sample t-test was conducted on the log-normalised data to test for an overall effect of predictability (attenuation of Repeat relative to Deviant sequences). Follow-up one-sample t-tests were then conducted separately for low and high stimulation intensities to determine whether predictability effects were present at each intensity level.

#### Experiments 2 and 3

A one-sample t-test was conducted on the log-normalised data to test for an overall effect of predictability (attenuation of global majority relative to global minority sequences). In Experiment 2, follow-up one-sample t-tests were conducted separately for the sequence type (Repeat, Deviant). In Experiment 3, follow-up one-sample t-tests examined predictability effects separately within each instruction condition (Informed, Uninformed).

## Supplemental Materials

### Raw MEP data

#### Experiment 1

Collapsed across expectation condition, high intensity stimulation produced significantly larger MEPs than low intensity stimulation, *F*(1,28) = 67.11, *p* <.001, η_p_^2^ = .706. Collapsed across stimulation intensity, MEPs elicited by repeat (expected) TMS did not significantly differ from MEPs elicited by deviant (unexpected) TMS, *F*(1,28) = 1.79, *p* = 0.191, η_p_^2^ = .060. However, a significant interaction between stimulation intensity and expectation condition, *F(*1,28) = 6.67, *p* = .015, η_p_^2^ = .192, suggested that prediction-based motor attenuation was numerically larger for high than low intensity stimulation.

#### Experiment 2

Collapsed across expectation condition, deviant sequences produced significantly larger MEPs than repeat sequences, *F*(1, 23) = 5.68, *p* = .026, η_p_^2^ = .198. Collapsed across sequence type, global majority (expected) sequences were weakly attenuated relative to global minority (unexpected) sequences, but this relationship was not statistically significant, *F*(1, 23) = 2.21, *p* = .151, η_p_^2^ = .087. A significant interaction between sequence type and expectation condition, *F*(1, 23) = 18.26, *p* < .001, η_p_^2^ = .443 (Figure 3A), indicated that the prediction-based attenuation effect was larger for deviant sequences than repeat sequences.

#### Experiment 3

Collapsed across information group, expected TMS produced significantly smaller MEPs than unexpected TMS, *F*(1, 67) = 4.75, *p* = .033, η_p_^2^ = .066. Collapsed across expectation condition, there was no significant difference in MEPs between the Informed and Uninformed group, *F*(1, 67) = 1.09, *p* = .300, η_p_^2^ = .016. Critically, a significant interaction between information group and expectation condition, *F*(1, 67) = 4.56, *p* = .036, η_p_^2^ = .064, indicated that attenuation differed as a function of explicit knowledge.

### Experiment 3 Post-Experimental Questionnaire

#### Screen 1

Each trial was indicated by a white fixation cross “+”. Did you notice any patterns or sequences to how the TMS was fired each trial? If so, please describe otherwise write “im not sure”.

TYPE your response immediately and then press ENTER to proceed to the next question.

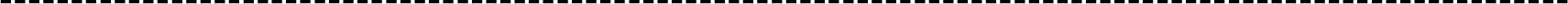

#### Screen 2

TMS always occurred in sequences of the same number of pulses. Did you notice how many TMS pulses made up a sequence?

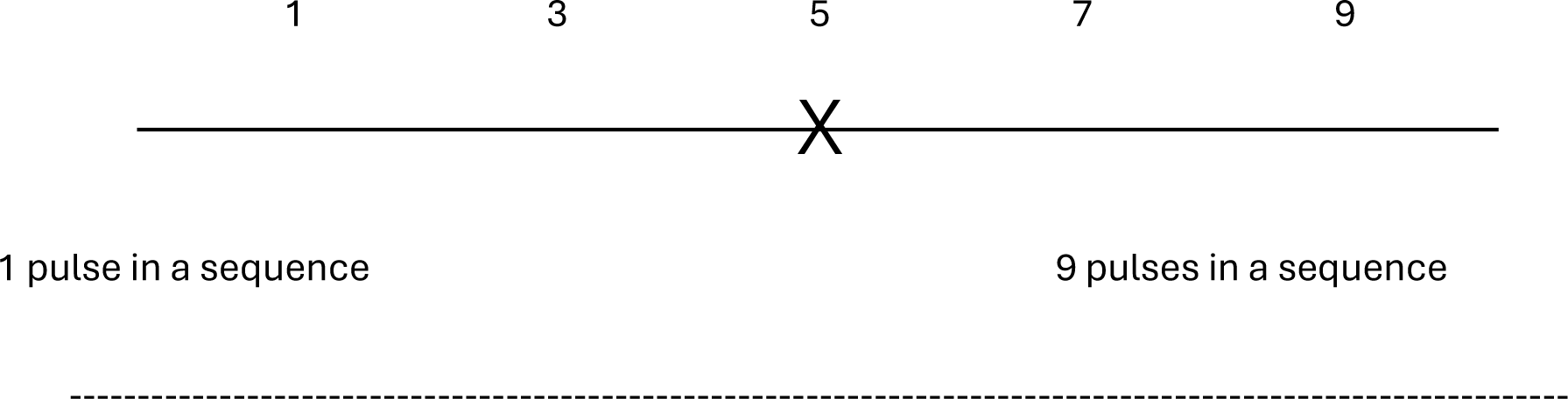

#### Screen 3

The TMS machine can be set to different intensities. This results in different “strengths” of TMS pulses and therefore different levels of activation of the motor system.

How many different intensities did you notice the TMS was fired at during the experiment?

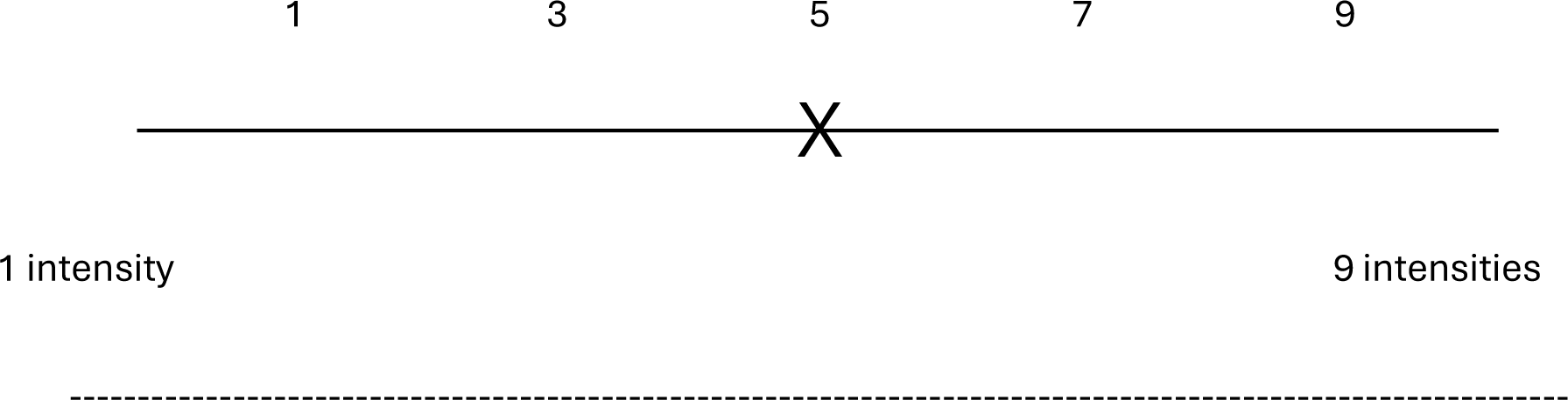

#### Screen 4

TMS pulses occurred in sequences of four (4) and there were two (2) different intensities (low and high) delivered during the experiment. Can you describe any sequence you noticed? For example, did you notice if they repeated or alternated between low and high in any way?

TYPE your response immediately and then press ENTER to proceed to the next question.

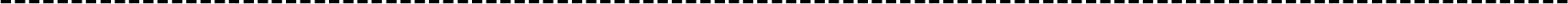

#### Screen 5

There were two (2) types of sequences:

Sequence 1: low, low, low, low
Sequence 2: low, low, low, high

In each block, did you notice a difference between the number of Sequence 1 and Sequence 2? Did this change across the blocks?

TYPE your response immediately and then press ENTER.

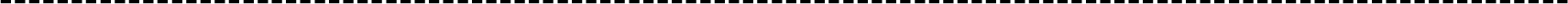

The post-experimental questionnaire comprised five questions combining free-recall and rating-scale formats, designed to assess participants’ explicit knowledge of the sequence structures and how their probabilities varied across blocks.

The first open-ended question failed to discriminate between participants with and without knowledge of the task structure. More critically, the third question, which asked participants to report how many distinct TMS pulse intensities were administered during the experiment, revealed a fundamental difficulty in using self-report to assess awareness in this paradigm.

Although the correct answer was two (low and high intensity), a substantial number of participants reported experiencing three or more distinct intensities, including participants in the Informed group who had been explicitly instructed that only two intensity levels existed. This finding suggests that the phenomenological experience of TMS stimulation is not straightforward to report, and that expectation may have modulated the subjective experience of stimulation intensity. Specifically, unexpected pulses of a given nominal intensity may have been perceived as subjectively different from expected pulses of the same intensity. This effect would be consistent with reports of expectation-dependent perceptual differences in other modalities (Dewey & Carr, 2013; Eagleman & Pariyadath, 2009). As a result, an incorrect response to this question does not necessarily indicate a lack of awareness of the experimental manipulations or sequence structure.

Removing the open-ended and intensity questions from scoring does not resolve issues of sensitivity. Variability in the subjective experience of stimulation intensity likely propagated to subsequent questions assessing knowledge of sequence types, as participants’ descriptions of the sequences would have been anchored to their perceptual experience of the pulses rather than their nominal intensity values. For these reasons, the questionnaire lacked the validity and sensitivity required to classify participants as aware or unaware of the predictive structure, and awareness classification was based solely on the instruction group manipulation.

## References

Akatsuka, K., Wasaka, T., Nakata, H., Kida, T., & Kakigi, R. (2007). The effect of stimulus probability on the somatosensory mismatch field. Experimental Brain Research, 181(4), 607–614. 10.1007/s00221-007-0958-4

Alilović, J., Slagter, H. A., & van Gaal, S. (2021). Subjective visibility report is facilitated by conscious predictions only. Consciousness and Cognition, 87, 103048. 10.1016/j.concog.2020.103048

Bekinschtein, T. A., Dehaene, S., Rohaut, B., Tadel, F., Cohen, L., & Naccache, L. (2009). Neural signature of the conscious processing of auditory regularities. Proceedings of the National Academy of Sciences - PNAS, 106(5), 1672–1677. 10.1073/pnas.0809667106

Clark, A. (2013). Whatever next? Predictive brains, situated agents, and the future of cognitive science. The Behavioral and brain sciences, 36(3), 181–204. 10.1017/S0140525X12000477

Clarke, J., Rittershofer, K., Ward, E. K., Yon, D., & Press, C. (2026). Perceptual predictions track subjective, over objective, statistical structure. eLife, 15. 10.7554/eLife.109690.1

den Ouden, H. E. M., Kok, P., & de Lange, F. P. (2012). How prediction errors shape perception, attention, and motivation. Frontiers in psychology, 3, 548–548. 10.3389/fpsyg.2012.00548

Dewey, J. A., & Carr, T. H. (2013). Predictable and self-initiated visual motion is judged to be slower than computer generated motion. Consciousness and cognition, 22(3), 987–995. 10.1016/j.concog.2013.06.007

Eagleman, D. M., & Pariyadath, V. (2009). Is subjective duration a signature of coding efficiency? Philos Trans R Soc Lond B Biol Sci, 364(1525), 1841–1851. 10.1098/rstb.2009.0026

Elijah, R. B., Le Pelley, M. E., & Whitford, T. J. (2018). Act Now, Play Later: Temporal Expectations Regarding the Onset of Self-initiated Sensations Can Be Modified with Behavioral Training. Journal of cognitive neuroscience, 30(8), 1145–1156. 10.1162/jocn_a_01269

Feuerriegel, D., Vogels, R., & Kovács, G. (2021). Evaluating the evidence for expectation suppression in the visual system. Neuroscience and biobehavioral reviews, 126, 368–381. 10.1016/j.neubiorev.2021.04.002

Fischer, C., Morlet, D., Bouchet, P., Luaute, J., Jourdan, C., & Salord, F. (1999). Mismatch negativity and late auditory evoked potentials in comatose patients. Clinical neurophysiology, 110(9), 1601–1610. 10.1016/s1388-2457(99)00131-5

Friston, K. (2005). A theory of cortical responses. Philosophical transactions of the Royal Society of London. Series B. Biological sciences, 360(1456), 815–836. 10.1098/rstb.2005.1622

Garrido, M. I., Kilner, J. M., Stephan, K. E., & Friston, K. J. (2009). The mismatch negativity: A review of underlying mechanisms. Clinical neurophysiology, 120(3), 453–463. 10.1016/j.clinph.2008.11.029

Hutchinson, J. B., & Barrett, L. F. (2019). The Power of Predictions: An Emerging Paradigm for Psychological Research. Current directions in psychological science: a journal of the American Psychological Society, 28(3), 280–291. 10.1177/0963721419831992

Knill, D. C., & Pouget, A. (2004). The Bayesian brain: the role of uncertainty in neural coding and computation. Trends in neurosciences (Regular ed.), 27(12), 712–719. 10.1016/j.tins.2004.10.007

Koelsch, S., Heinke, W., Sammler, D., & Olthoff, D. (2006). Auditory processing during deep propofol sedation and recovery from unconsciousness. Clinical neurophysiology, 117(8), 1746–1759. 10.1016/j.clinph.2006.05.009

Koshiyama, D., Kirihara, K., Tada, M., Nagai, T., Fujioka, M., Usui, K., Araki, T., & Kasai, K. (2020). Reduced Auditory Mismatch Negativity Reflects Impaired Deviance Detection in Schizophrenia. Schizophrenia bulletin, 46(4), 937–946. 10.1093/schbul/sbaa006

Lee, T. S., & Mumford, D. (2003). Hierarchical Bayesian inference in the visual cortex. Journal of the Optical Society of America. A, Optics, image science, and vision, 20(7), 1434–1448. 10.1364/josaa.20.001434

Liang, L., Tam, C. K. W., Livesey, E. J., & Tran, D. M. D. (2025). Awareness is necessary for predictive learning and prediction-based motor attenuation. bioRxiv, 2025.2012.2001.691731. 10.64898/2025.12.01.691731

Lieder, F., Daunizeau, J., Garrido, M. I., Friston, K. J., & Stephan, K. E. (2013). Modelling trial-by-trial changes in the mismatch negativity. PLoS computational biology, 9(2), e1002911–e1002911. 10.1371/journal.pcbi.1002911

Meijs, E. L., Slagter, H. A., de Lange, F. P., & van Gaal, S. (2018). Dynamic interactions between top-down expectations and conscious awareness. Journal of Neuroscience, 38(9), 2318–2327. 10.1523/JNEUROSCI.1952-17.2017

Naccache, L., Puybasset, L., Gaillard, R., Serve, E., & Willer, J.-C. (2005). Auditory mismatch negativity is a good predictor of awakening in comatose patients: a fast and reliable procedure. Clinical neurophysiology, 116(4), 988–989. 10.1016/j.clinph.2004.10.009

Normann, R. A., & Perlman, I. (1979). The effects of background illumination on the photoresponses of red and green cones. The Journal of physiology, 286(1), 491–507. 10.1113/jphysiol.1979.sp012633

Nourski, K. V., Steinschneider, M., Rhone, A. E., Kawasaki, H., Howard, R. M. A., & Banks, M. I. (2018). Auditory Predictive Coding across Awareness States under Anesthesia: An Intracranial Electrophysiology Study. The Journal of neuroscience, 38(39), 8441–8452. 10.1523/JNEUROSCI.0967-18.2018

Pfeiffer, C., Nguissi, N. A. N., Chytiris, M., Bidlingmeyer, P., Haenggi, M., Kurmann, R., Zubler, F., Accolla, E., Viceic, D., Rusca, M., Oddo, M., Rossetti, A. O., & De Lucia, M. (2018). Somatosensory and auditory deviance detection for outcome prediction during postanoxic coma. Annals of clinical and translational neurology, 5(9), 1016–1024. 10.1002/acn3.600

Rao, R. P. N., & Ballard, D. H. (1999). Predictive coding in the visual cortex: a functional interpretation of some extra-classical receptive-field effects. Nature neuroscience, 2(1), 79–87. 10.1038/4580

Rideaux, R., Dang, P., Jackel-David, L., Buhmann, Z., Rangelov, D., & Mattingley, J. B. (2025). Violated predictions enhance the representational fidelity of visual features in perception. Journal of Vision, 25(4), 14–14. 10.1167/jov.25.4.14

Rossi, S., Antal, A., Bestmann, S., Bikson, M., Brewer, C., Brockmöller, J., Carpenter, L. L., Cincotta, M., Chen, R., Daskalakis, J. D., Di Lazzaro, V., Fox, M. D., George, M. S., Gilbert, D., Kimiskidis, V. K., Koch, G., Ilmoniemi, R. J., Lefaucheur, J. P., Leocani, L.,…Hallett, M. (2021). Safety and recommendations for TMS use in healthy subjects and patient populations, with updates on training, ethical and regulatory issues: Expert Guidelines. Clinical neurophysiology, 132(1), 269–306. 10.1016/j.clinph.2020.10.003

Rossini, P. M., Burke, D., Chen, R., Cohen, L. G., Daskalakis, Z., Di Iorio, R., Di Lazzaro, V., Ferreri, F., Fitzgerald, P. B., George, M. S., Hallett, M., Lefaucheur, J. P., Langguth, B., Matsumoto, H., Miniussi, C., Nitsche, M. A., Pascual-Leone, A., Paulus, W., Rossi, S.,…Ziemann, U. (2015). Non-invasive electrical and magnetic stimulation of the brain, spinal cord, roots and peripheral nerves: Basic principles and procedures for routine clinical and research application. An updated report from an I.F.C.N. Committee. Clinical neurophysiology, 126(6), 1071–1107. 10.1016/j.clinph.2015.02.001

Schnakers, C., & Laureys, S. (2012). Coma and Disorders of Consciousness. Springer.

Schultz, W., Dayan, P., & Montague, P. R. (1997). A neural substrate of prediction and reward. Science, 275(5306), 1593–1599. 10.1126/science.275.5306.1593

Sepulcre, J., Sabuncu, M. R., Yeo, T. B., Liu, H., & Johnson, K. A. (2012). Stepwise connectivity of the modal cortex reveals the multimodal organization of the human brain. Journal of Neuroscience, 32(31), 10649–10661. 10.1523/jneurosci.0759-12.2012

Simpson, T. P., Manara, A. R., Kane, N. M., Barton, R. L., Rowlands, C. A., & Butler, S. R. (2002). Effect of propofol anaesthesia on the event-related potential mismatch negativity and the auditory-evoked potential N1. British journal of anaesthesia: BJA, 89(3), 382–388. 10.1093/bja/89.3.382

Sriutaisuk, N., & Franz, E. A. (2025). Predictable transcranial magnetic stimulation suppresses corticospinal excitability: a TMS experiment. Experimental Brain Research, 243(6), 134. 10.1007/s00221-025-07091-y

Stevens, J. C., & Stevens, S. S. (1963). Brightness function: effects of adaptation. Journal of the Optical Society of America (1930), 53(3), 375-385. 10.1364/JOSA.53.000375

Strauss, M., Sitt, J. D., King, J.-R., Elbaz, M., Azizi, L., Buiatti, M., Naccache, L., van Wassenhove, V., & Dehaene, S. (2015). Disruption of hierarchical predictive coding during sleep. Proceedings of the National Academy of Sciences - PNAS, 112(11), E1353–E1362. 10.1073/pnas.1501026112

Summerfield, C., Trittschuh, E. H., Monti, J. M., Mesulam, M. M., & Egner, T. (2008). Neural repetition suppression reflects fulfilled perceptual expectations. Nat Neurosci, 11(9), 1004–1006. 10.1038/nn.2163

Tang, M. F., Smout, C. A., Arabzadeh, E., & Mattingley, J. B. (2018). Prediction error and repetition suppression have distinct effects on neural representations of visual information. eLife, 7. 10.7554/eLife.33123

Tivadar, R. I., Knight, R. T., & Tzovara, A. (2021). Automatic Sensory Predictions: A Review of Predictive Mechanisms in the Brain and Their Link to Conscious Processing. Frontiers in human neuroscience, 15, 702520–702520. 10.3389/fnhum.2021.702520

Tran, D. M. D., & Livesey, E. J. (2021). Prediction-based attenuation as a general property of learning in neural circuits. Journal of Experimental Psychology: Animal Learning and Cognition, 47(1), 14. 10.1037/xan0000280

Tran, D. M. D., McNair, N. A., Harris, J. A., & Livesey, E. J. (2021). Expected TMS excites the motor system less effectively than unexpected stimulation. NeuroImage, 226, 117541–117541. 10.1016/j.neuroimage.2020.117541

Tran, D. M. D., McNair, N. A., Whitton, A. E., Whitford, T. J., & Livesey, E. J. (2025). Prediction-based sensory attenuation is related to prediction-based motor attenuation. Imaging Neuroscience, 3, *IMAG-a.* 10.1162/IMAG.a.1045

Tschentscher, N., Woll, C. F. J., Tafelmaier, J. C., Kriesche, D., Bucher, J. C., Engel, R. R., & Karch, S. (2023). Neurocognitive Deficits in First-Episode and Chronic Psychotic Disorders: A Systematic Review from 2009 to 2022. Brain Sciences, 13(2), 299. https://www.mdpi.com/2076-3425/13/2/299

Tzovara, A., Fedele, T., Sarnthein, J., Ledergerber, D., Lin, J. J., & Knight, R. T. (2024). Predictable and unpredictable deviance detection in the human hippocampus and amygdala. Cereb Cortex, 34(2). 10.1093/cercor/bhad532

van Schalkwyk, G. I., Volkmar, F. R., & Corlett, P. R. (2017). A Predictive Coding Account of Psychotic Symptoms in Autism Spectrum Disorder. J Autism Dev Disord, 47(5), 1323–1340. 10.1007/s10803-017-3065-9

Weidemann, G., Satkunarajah, M., & Lovibond, P. F. (2016). I Think, Therefore Eyeblink: The Importance of Contingency Awareness in Conditioning. Psychological science, 27(4), 467–475. 10.1177/0956797615625973

